# Designing epitope-based vaccines against Nipah virus Glyco and Fusion proteins using integrated immunoinformatics and structural modeling techniques

**DOI:** 10.64898/2026.01.07.698100

**Authors:** Sattyajit Biswas, Md. Kamaruzzaman, Minhaj Bin Sayem Alam, Md. Jafor Iqbal, Maryam Jamila, Tanzil Zahan

## Abstract

Nipah virus (NiV) is a re-emerging zoonotic virus belonging to the Paramyxoviridae family that results in significant neurological damage and raises fatality rates. NiV is classified as a stage III pathogen with a high likelihood of transmission to humans, causing outbreaks intermittently and without any predictable pattern. In Bangladesh, outbreaks of NiV have happened every year since 2001. Even though the disease is severe, no antiviral medications exist for NiV infections. As a consequence, developing a vaccine is crucial. The present research aims to predict an effective epitope-based vaccine by applying immunoinformatic techniques against the fusion and glycoprotein of the Nipah virus. Fusion and glycoproteins were obtained from the UniProt protein database and screened for the T and B cell epitopes using the IEDB and ABCpred servers. Moreover, the constructed 3D structure of the NiV vaccine was occupied with Toll-like receptor 4 for molecular docking and dynamic (MD) simulation studies. Finally, the vaccine design was validated for expression in the PET28a (+) vector of *Escherichia coli,* and immunological simulations were also conducted. After determination of the allergenicity, antigenicity, and toxicity, the non-allergenic, nontoxic, antigenic, and immunogenic epitopes were used to construct the vaccine with adjuvants and an appropriate linker, including AAY, GPGPG, and KK. The constructed NiV vaccine was confirmed based on its physicochemical properties and docking scores for further analysis. The MD simulations indicated a stabilized structure and increased duration of epitope visibility, indicating strong immune responses. Furthermore, codon optimization and in silico cloning were used to verify bacterial expression of the developed NiV vaccine in *E. coli*. The results showed that NiV can increase immune responses against the Nipah virus. Furthermore, *in vitro* and *in vivo* research and clinical studies are recommended to establish the findings of this study.

## Introduction

Nipah virus (NiV) is a highly pathogenic RNA virus with a negative-sense genome that belongs to the Paramyxoviridae family. Fruit bats are the most common hosts of this virus. It can spread to intermediate hosts like pigs and humans, proving its zoonotic potential across multiple species (1), (2). The human Nipah virus (NiV) affected 276 individuals in Malaysia and Singapore during the initial epidemic, which occurred throughout 1998 and 1999, along with the reported mortality ratio for the documented cases ranging from 40% to 100%, revealing NiV as a severe hazard to public health issues (3,4), (5). Since then, several outbreaks were reported in different countries, especially in South and Southeast Asia, such as in India, Bangladesh, and the Philippines. These areas are highly concentrated with infections, and they are experiencing an outbreak of the infection continuing to the present day (6), (7), (8),(9). There are two genetically separate strains of the Nipah virus (NiV): the Malaysian strain (NiV M) and the Bangladeshi strain (NiV B). The variation in the dynamics of the virus distribution and death rate also indicates that NiV B has a greater pathogenicity than NiV M (10). As a result, the World Health Organization (WHO) R&D Blueprint does treat it as a high-priority pathogen concerning the priorities of critical research and development (11). The signs of Nipah virus (NiV) infection may include muscle pains and flu-like symptoms such as fever, coughing, nausea, light-headedness, and headaches. In later stages, it may evolve into significant health issues, among them acute inflammation of the brain (encephalitis), inflammation of blood vessels in general (systemic vasculitis), and difficulty breathing (12), (1), (13). It is categorized as a Biosafety level 4 pathogen based on its wide host range with high virulence that has the potential to cause severe disease and mortality (14).

NiV has a genome composed of six structural proteins in a single-stranded non-segmented RNA approximately 18.2 kilobases in size, namely: a matrix protein (M), a fusion protein (F), a phosphoprotein (P), an RNA polymerase (L), a glycoprotein (G), and a nucleocapsid (N) (15). Among them, G and F proteins are the two main envelope glycoproteins that connect to the virus and allow it to attach to the host cell (16). The N-linked glycans on the G protein enable the virus’s initial attachment to the host cell, while the F protein plays an important role in the fusion process between the host cell membrane and the viral envelope (16), (17). The virus enters through class B2/B3 Ephrin receptors, which are mostly found in the respiratory tract and the vascular tissue. This interaction triggers the structural changes that the G protein mediates, which in turn activate the F glycoprotein to complete the membrane fusion (18), (19). Although NiV is an alarming new illness with the potential to spread globally, there are presently no approved treatments or vaccinations for use in humans or any other intermediate hosts of the virus (20). G and F proteins are also the main targets of antibody neutralization and are considered the key reagents used in vaccine-based immunity to this virus (21).

Over the last few decades, bioinformatics and structural biology have experienced revolutionary development. The subsequent progress of the computational methods of analyzing genomic data has also led to the creation of new methods for the production of vaccines. New research focuses on implementing immunoinformatics-based strategies to engineer next-generation multi-epitope vaccines, which become practical in the sense that they avoid labor-intensive pathogen cultivation and dramatically speed up the period of vaccine development (22). By evaluating the pathogen’s proteome profile using bioinformatic tools and computational analysis, researchers will have access to discover accessible epitopes to trigger an immune response (23). These newly identified epitopes may be constructed into a multi-epitope subunit vaccination, which offers various advantages, such as lower risks of viral outbreaks, increased immune response, and greater durability compared to normal virus-based vaccines (24). This novel technique has significant implications for clinical research and can successfully combat viral infections (25). Our present study aims to create a multi-epitope-based peptide vaccination against the Nipah virus, incorporating the G and F proteins. This method builds on current understanding of the immunological responses elicited by other protein members of the paramyxovirus group (26), (27) and evaluates its efficacy using computational and immunoinformatics methods.

## Methodology

A summary of the systematic approach utilized in this work to develop and evaluate the NiV multi-epitope vaccine candidate is presented in **Fig 1**.

**Fig 1.**
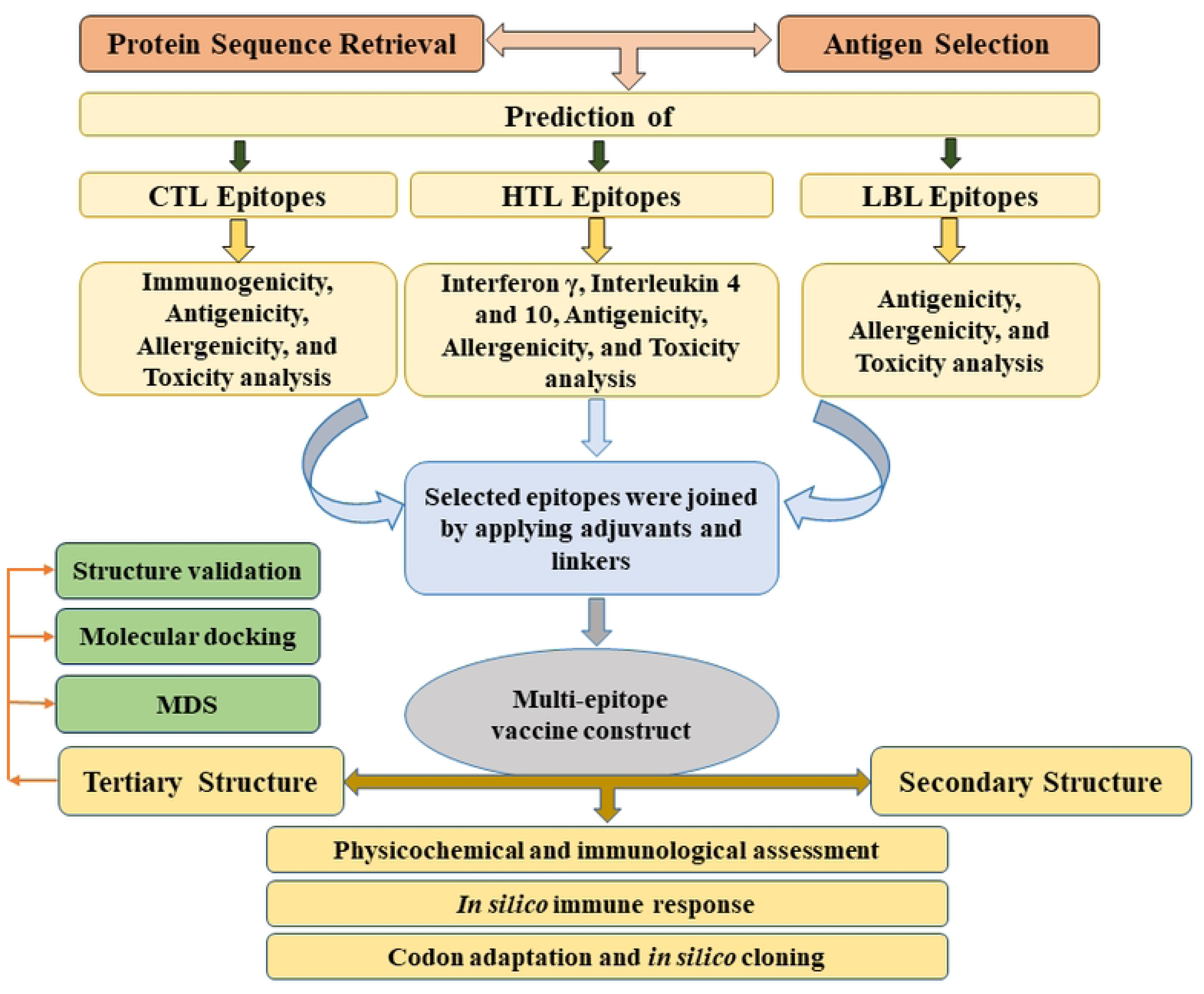
**S**chematic illustration of the complete workflow for designing a multi-epitope vaccine from Nipah virus fusion and glycoproteins.

### Retrieval of protein sequence and estimating antigenicity

During the retrieval of the protein sequence, the F and G glycoproteins of the Nipah virus were obtained from the UniProt database (https://www.uniprot.org/). Following that, the detected protein sequences were submitted to the sequence identity and similarity (SIAS) database to determine the similarity of all protein sequences (http://imed.med.ucm.es/Tools/sias.html). These sequences were subsequently examined to assess the antigenic potential of the proteins using the VaxiJen v2.0 server (https://www.ddg-pharmfac.net/vaxijen/VaxiJen/VaxiJen.html) (28) with a threshold value of 0.4, which has been used to screen viral proteins (29). Every protein with a score higher than this threshold was considered antigenic, while any protein with a value lower than 0.4 was described as non-antigenic.

### Prediction of T-cell and B-cell immunogenic epitopes

The IEDB site (https://www.iedb.org/) was utilized to screen MHC class I (CTL) and MHC class II (HTL) epitopes in the structural proteins G and F of Nipah virus (NiV). It was predicted through NetMHCIIpan 4.1 EL tools online to give in vivo and in vitro (predicted) ranking among epitopes and epitopes of cytotoxic and helper T cells (30). The CTL epitope search was done with 9 to 10-mer peptides, and HTL epitope prediction with 15-mer peptides, with the default reference allele set applied. To validate the antigenic property of the selected epitopes, the VaxiJen version 2.0 was used, and the threshold value of 0.4 was applied to predict the candidates (29). In parallel, the MHC I immunogenicity tool (http://tools.iedb.org/immunogenicity/) through the IEDB portal was also utilized to predict the immunogenicity of the CTL epitopes. Epitopes with a positive immunogenicity score only were shortlisted for further evaluation (31). Conversely, HTL epitopes were evaluated in terms of their ability to trigger production of IFN-gamma, an important component of priming the innate and adaptive immune responses against viral infections. This study was also carried out using the IFNepitope server (https://webs.iiitd.edu.in/raghava/ifnepitope/predict.php). All HTL epitopes with a positive reaction in IFN-gamma were used in further research (32). To identify probable B-cell epitopes of the NiV proteins, the web-based B-cell epitope prediction program ABCpred (https://webs.iiitd.edu.in/raghava/abcpred/ABC_submission.html) was used. In epitope prediction, where only 16-mer peptides were considered, the selection threshold exceeded 0.50 (33). Since the epitopes that were predicted could be used to perform antigenicity analysis, a server known as VaxiJen v2.0 (29) was utilized to perform the analysis

### Population coverage of predicted epitopes

The frequency of particular HLA alleles between various ethnic groups and populations is necessary for developing an advantageous epitope-based vaccine. The IEDB population coverage tool (http://tools.iedb.org/population/) was used to determine an assessment of the representations of the selected CTL and HTL epitopes across these alleles. The tool projected a cumulative coverage level of HLA class I and II alleles at the regional and global levels (34).

### Identification of immunogenic epitopes and vaccine construction

T and B cell epitopes are crucial in the initial immune responses and play a key role in developing multiepitope vaccines. This approach uses various data processing techniques to create a vaccine that includes multiple pathogen identification tests by targeting a combination of T and B cell epitopes. From the IEDB and ABCpred server, T and B cell epitopes were further evaluated using ToxinPred 2.0 (35), AllerTOP v2.0 (28), and Vaxijen 2.0 (29) to identify the final epitopes for vaccine construction. To design a multi-epitope vaccine, the epitopes of cytotoxic T cells and helper T cells are conjugated with Type AAY and GPGPG, respectively, and those of B cells are attached with KK (36,37). Enhance the immunity, conjugating the adjuvant with the epitopes was done, and beta defensin 3 (Accession ID: P81534) was selected as the adjuvant to develop the vaccine. Epitopes were linked to the adjuvant through several linkers, to begin with the EAAAK linker. The EAAAK linker can be added to the adjuvant at the vaccine’s N terminus to increase its immunogenicity (38). In multiepitope vaccinations, linkers have the correct spacing of various epitopes, thus preserving structural integrity, allowing each to elicit its immune response without interference.

### Physicochemical studies of the constructed vaccine

Detailed analysis of the physicochemical characteristics is a vital step in vaccine development that must pass adequate scrutiny following its effectiveness, safety, and compatibility with the human organic system. To this end, ExPASy ProtParam (https://web.expasy.org/protparam/) was used to analyze the major physiological parameters of the developed vaccine. Utilizing this platform allowed for the identification and detailed examination of several essential properties concerning the vaccine construct, such as aliphatic index, stability index, Grand Average of Hydropathicity (GRAVY), and half-life (39). The SoluProt 1.0 analysis tool, found at https://loschmidt.chemi.muni.cz/soluprot/, was used to determine the vaccine’s solubility (40).

### Predicting the 2D and 3D structure of the vaccine

The PSIPRED tool (http://bioinf.cs.ucl.ac.uk/psipred/) was used to predict the secondary structural components, including alpha helices, beta sheets, and coils, present in the vaccine constructs (41). The GalaxyTBM server (https://galaxy.seoklab.org/cgi-bin/submit.cgi?type=TBM) was employed in modeling the three-dimensional structure of the multi-epitope, multi-target vaccine peptide. This server builds a solid backbone framework based on various structure templates and completes uncertainties by ab initio modeling of loop or termini to the rigid framework established (42).

### Optimizing and validating the vaccine’s 3D structure

To increase the structural accuracy of the predicted 3D model, refinement was done through the Galaxy Refine server (http://galaxy.seoklab.org/refine) (43). The effectiveness and durability of the constructed vaccination models were afterward confirmed by structural evaluation servers such as ERRAT and PROCHECK (44) consolidated within the SAVES v6.0 program (https://saves.mbi.ucla.edu/) and PROSA-Web server (https://prosa.services.came.sbg.ac.at/prosa.php) (45).

### Identification of conformational B-cell epitopes

Conformational B-cell epitopes were tested upon prediction and correction of the 3D structure of the final vaccine candidate to identify epitopes that could generate B-cell responses. To achieve this, the ElliPro tool on the IEDB database (http://tools.iedb.org/ellipro/), which assesses the linearity of the protein’s structure or sequence to identify regions with discontinuous inheritance of antibodies (46).

### *In-silico* molecular docking analysis with the vaccine’s 3D structure and TLR-4 receptor

Molecular docking analysis revealed the interaction between a ligand and its receptor, predicting the energy score produced during that contact. Using specific grading systems, this approach can ascertain the binding affinities of two compounds (47). The target Toll-like receptor 4 (TLR4), with PDB ID 4G8A, was obtained from the RCSB Protein Data Bank (https://www.rcsb.org/) for molecular docking (48). The vaccine candidates identified as ligands were docked with the TLR4 receptor. The ClusPro 2.0 web server (https://cluspro.org/login.php) was used for molecular docking to assess the binding affinity. It studies the three-dimensional structure of proteins, predicts areas where ligands bind, and reveals sites that may have interactions between proteins and small molecules. The best-docked complex was chosen and recovered based on the ligand-receptor complex’s binding affinity (49).

### Molecular dynamics simulation

To get the consistency of the expected vaccine and vaccine receptor (VR) complex, a computational molecular dynamics (MD) simulation of the enhanced vaccine and VR complex was performed using the Linux operating system and the ‘Desmond v6.3 Program’ in the Schrodinger (academic version). This computational approach has been applied to obtain the thermodynamic stability of the vaccination and VR combination, simulating water with OPLS3e as the force field and the pre-constructed TIP3P water model (50). Rectangular constraints on the border representing the periodical structure were made to define the size and shape of the unit repeating, which is being shielded by 20 Angstroms spacings. An electrical neutralization was obtained by balancing proper sodium and chlorine ions to achieve a minimum charge within the Desmond module. This was done through the OPLS3eforce field. Molecular dynamics simulations were run with periodic boundaries in constant particles, pressure, and temperature (NPT) ensemble (51). After this, the system architecture was optimized to utilize a protocol that considers a protocol with OPLS3e force field parameters, in the Desmond simulation suite of products. NPT ensembles were controlled at 300 K temperature and 1.01325 bar pressure using Nose-Hoover thermostat coupling and an isotropic pressure control technique to maintain stability. Every simulation set implied 50 PS relaxation progresses and a restraint force of 1.2 kcal/mol (51,52). The stability of the vaccination complex was further measured using the root mean square deviation (RMSD), root mean square fluctuation (RMSF), and radius of gyration (rGyr).

### Disulfide bond optimization, host-specific codon adaptation, and in-silico cloning

The Disulfide by Design 2.12 web server (http://cptweb.cpt.wayne.edu/DbD2/index.php) was used in Disulfide engineering, which is an experimental and broadly used technique introduced to create disulfide bonds to improve the structural stability of the vaccine construct (53). After structural refinement, the identified vaccine construct was codon optimized and subjected to in silico cloning. To determine the potentiality of the designed multi-epitope vaccine to be expressed in *E. coli* strain K12, the codon optimization tool offered by VectorBuilder (https://en.vectorbuilder.com/tool/codon-optimization) was utilized to characterize the possible expression efficiency. The tool examined the submitted sequence region on its GC content and Codon Adaptation Index (CAI) grounds to make it compatible with the host system and give it higher expression. Ideal CAI is closer to 1.0, and GC content between 30-70% is optimal for effective transcription and translation (54). Lastly, Snap Gene software (https://www.snapgene.com/) was used to design the insertion of the optimized vaccine construct into the pET-28a (+) expression vector, ensuring that the vector is correctly designed and compatible.

### In silico simulation of immunological responses

The designed vaccine was simulated via its immunogenic potential with immunological simulations by the C-ImmSim platform (https://kraken.iac.rm.cnr.it/C-IMMSIM/index.php). This simulation program uses a position-specific score matrix (PSSM) and various algorithmic methods to predict and analyse epitope and immune interactions. The simulation structure was set to simulate a normal vaccination regime, with a four-week delay between doses. Injections used during the four weeks were administered at time steps 1, 84, and 168, designating three simulated injections. The simulation volume was originally set to 10, and immune response analysis was performed throughout 1,000 simulation steps (55).

## Results

### Protein sequence retrieval and primary screening of the sequences

Initially, 50 protein sequences were obtained from the UniProt database. To prevent redundancy, similarity evaluation of the protein sequences was performed using the sequence identity and similarity (SIAS) server. The most similar proteins were removed from the selection list (**S1 Table)**. After that, the antigenic potential of the non-redundant proteins was assessed on the VaxiJen v2.0 server. Moreover, allergenicity and toxicity profiling were conducted to ensure the development of safe and immunogenic candidates. Finally, two antigenic, non-allergenic, and non-toxic Glycoproteins (ID: A0A7L5MR85 and A0A7L5MRB2) and fusion proteins (ID: A0A1L7B8D7 and A0A7M4C6A7) were chosen for further analysis, as displayed in **Table 1**.

**Table 1:** Nipah virus structural proteins retrieved along with their accession IDs and predicted VaxiJen v2.0 scores.

| Nipah virus structural protein | Accession ID | Antigenicity Score | Prediction |
| --- | --- | --- | --- |
| Fusion protein | A0A1L7B8D7 | 0.5012 | Probable Antigen |
|  | A0A7M4C6A7 | 0.5160 | Probable Antigen |
| Glycoprotein | A0A7L5MR85 | 0.4893 | Probable Antigen |
|  | A0A7L5MRB2 | 0.5083 | Probable Antigen |

### Detection of T-cell and B-cell responsive epitopes

All structural proteins were analysed to predict T-cell epitopes (HTL and CTL) using the IEDB server with NetMHCpan 4.1 EL and NetMHCIIpan 4.1 EL, using a full set of HLA reference to reach a global coverage of common MHC class I and II alleles.

NetMHCpan 4.1 EL detected 778 and 756 epitopes in Glyco and Fusion proteins, respectively, whereas 1312 and 1273 HTL epitopes were predicted on Glyco and Fusion proteins by NetMHCIIpan 4.1 EL. Refinement of candidate epitopes was established based on allele interaction frequency, antigenicity, non-toxicity, strain-wide conservation, and non-allergenicity. The immunogenicity of CTL epitopes was also measured using the IEDB MHC class I tool, and the IFN-assembly induction potential of HTL epitopes was measured through the IFN-epitope server. The prominent epitopes of each protein were included; 4 CTL epitopes were included in the final vaccine construct out of a total of 13 analyzed in CTL epitopes, and 3 HTL epitopes out of a total of 5 analyzed in HTL epitopes (**Table 2 and S2-S3 Table**). ABCpred was used to predict the linear B-cell epitopes of the designed protein. In the case of four proteins, 16 possible LBL epitopes were considered highly antigenic (> 0.5) and assessed as non-allergenic and non-toxic. According to these criteria, 5 epitopes were shortlisted to be considered further. **Table 2 and S4 Table** show the predicted epitopes with their starting and ending positions, magnitude, and scores.

**Table 2:** Selected CTL, HTL, and LBL epitopes with relevant properties such as antigenicity score, allergenicity, and toxicity for multiepitope vaccine design.

| MHC-I Epitopes | Antigenicity | Immuno-<br>genicity | MHC-II Epitopes | Antigeni-<br>city | Interfer-<br>on-γ | B-Cell Epitopes | Score | Allerge-<br>nicity | Toxic<br>ity |
| --- | --- | --- | --- | --- | --- | --- | --- | --- | --- |
| SIVPNFILV | 0.5759 | 0.21198 | TFISFIIVE<br>KKRNTY | 1.4822 | Positive | LGSVNYNSEGIAIGPP | 0.86 | Non<br>allergen | Non<br>toxin |
| ITFISFIIV | 1.3332 | 0.3111 | TEIQQAYI<br>QELLPVS | 0.5068 | Positive | SFIIVEKKRNTYSRLE | 0.65 | Non<br>allergen | Non<br>toxin |
| TLYFPAVGF | 0.7663 | 0.19727 | PAVGFLV<br>RTEFKYN<br>D | 1.4926 | Positive | SSTITIPANIGLLGSK | 0.67 | Non<br>allergen | Non<br>toxin |
| AMDEGYFA<br>Y | 0.7221 | 0.25913 |  |  |  | PLLAMDEGYFAYSHL<br>E | 0.7 | Non<br>allergen | Non<br>toxin |
|  |  |  |  |  |  | YRAQLASEDTNAQKTI | 0.8 | Non<br>allergen | Non<br>toxin |

### Estimation of population coverage based on HLA allele frequencies

The vaccine construct targeted a wide human ethnic coverage. To accomplish these goals, the IEDB population coverage database establishes the worldwide coverage of the projected vaccine epitopes. The T-cell epitope predictors consist of 22 HLA alleles that cover 97.94% of the human population. The comprehensive population coverage of the vaccine is shown in **Fig 2**. These results suggest that the vaccine epitopes can induce immune responses in the broad majority of people in the world. Moreover, the population coverage analysis revealed a heavy prevalence in areas that had past Nipah outbreaks, especially South and Southeast Asia. The population statistics of these regions are derived from the world population, indicating the estimated extent of coverage for India, Bangladesh, Malaysia, Singapore, and the Philippines, where the incidence of Nipah virus outbreaks is common.

**Figure 2:**
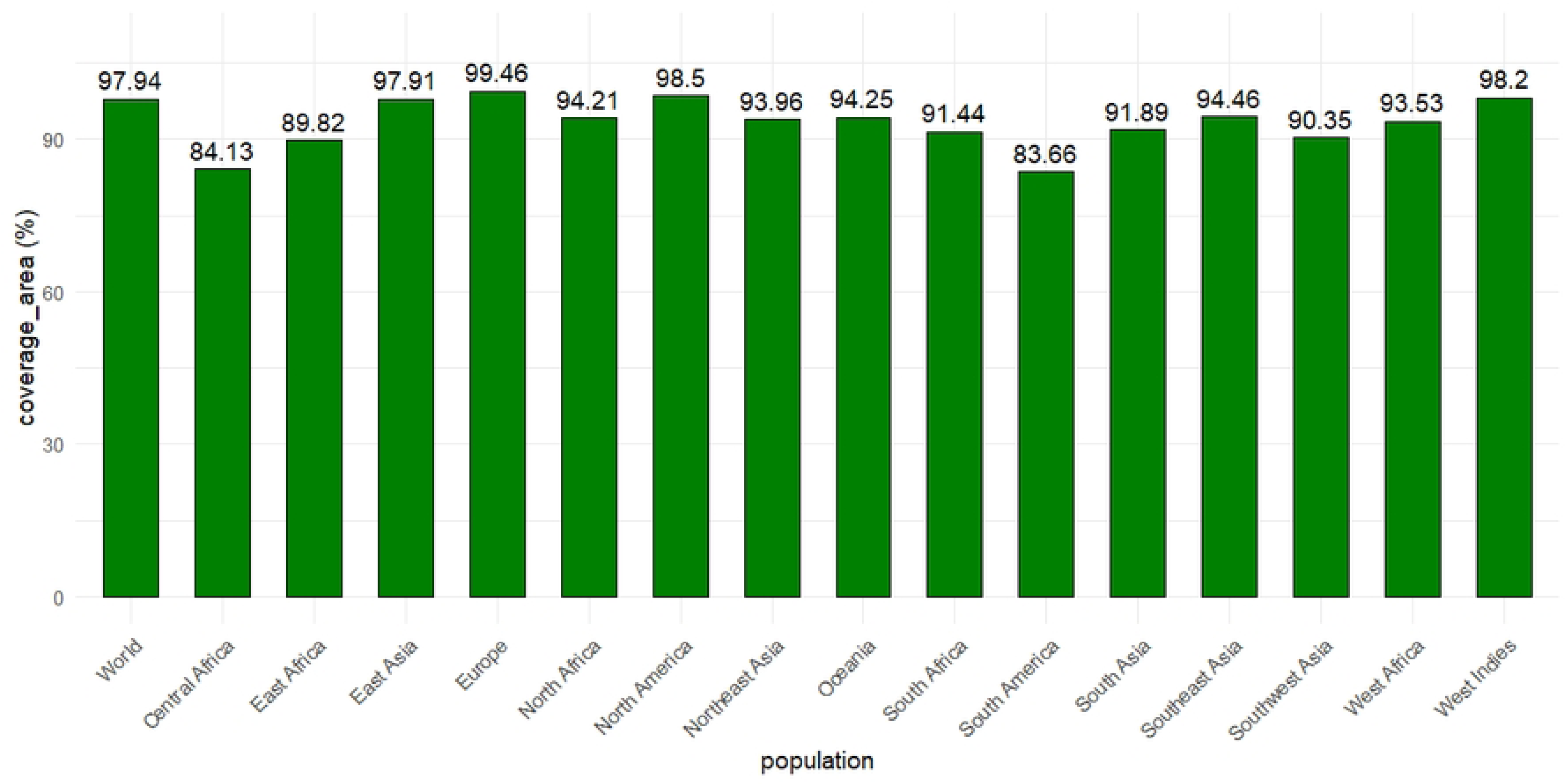
Worldwide population coverage analysis with the HLA alleles and the selected epitopes. The green colour indicates the percentage of T cell epitopes.

### Creation and structural design of a multi-target epitope vaccine construct

To combine antigenic epitopes into a single, unique structure, linkers and adjuvants such as Beta defensin 3 were employed. Due to its strong immunomodulating and antivirus effects, the ability to stimulate immune response, and the effectiveness of vaccination, beta-defensin has been selected by this work as an adjuvant in formulating the Nipah virus vaccine. Common designs were preferred in the investigation due to the problem of generating numerous potential epitope configurations. These structures were in different orders depending on the adjuvant and epitope arrangement. These constructs containing 4 CTL, 5 B-cell, and 3 HTL epitopes, excluding overlaps and similarity, were concatenated with universal linkers (EAAAK, AAY, KK, and GPGPG) as demonstrated in **Fig 3**.

**Fig 3.**
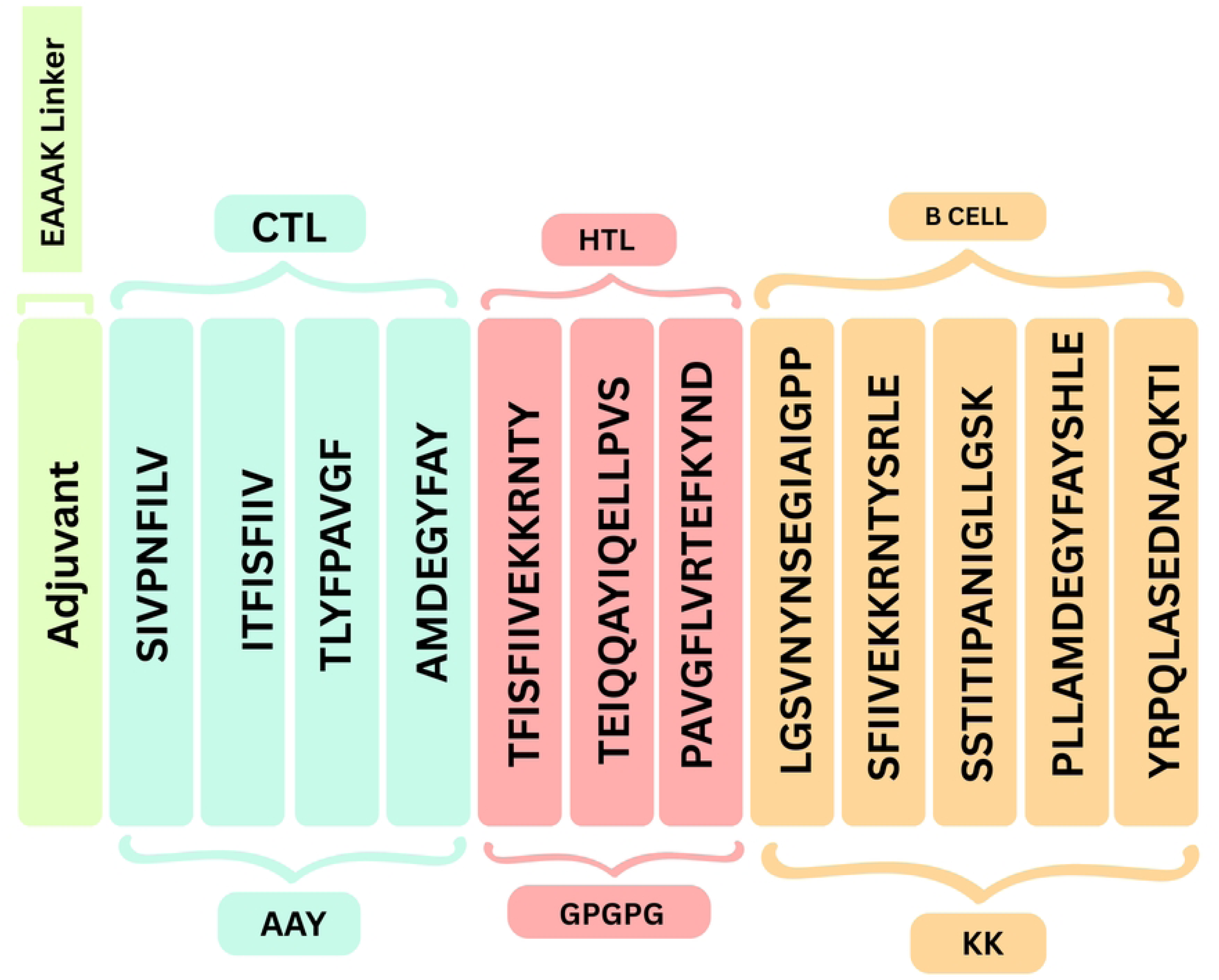
Graphical illustration of the multi-epitope vaccine construction. β-defensin 3 adjuvant was attached at the N-terminal, and CTL, HTL, and B cell epitopes were linked with the AAY, GPGPG, and KK linkers to form the vaccine.

### Analysis of physicochemical properties of the vaccine

Assessment of the physicochemical parameters was performed using metrics such as the molecular weight, amino acid count, theoretical pI, the aliphatic index (AI), grand average of hydropathicity (GRAVY), estimated half-life, and instability index (**Table 3**).

**Table 3:**
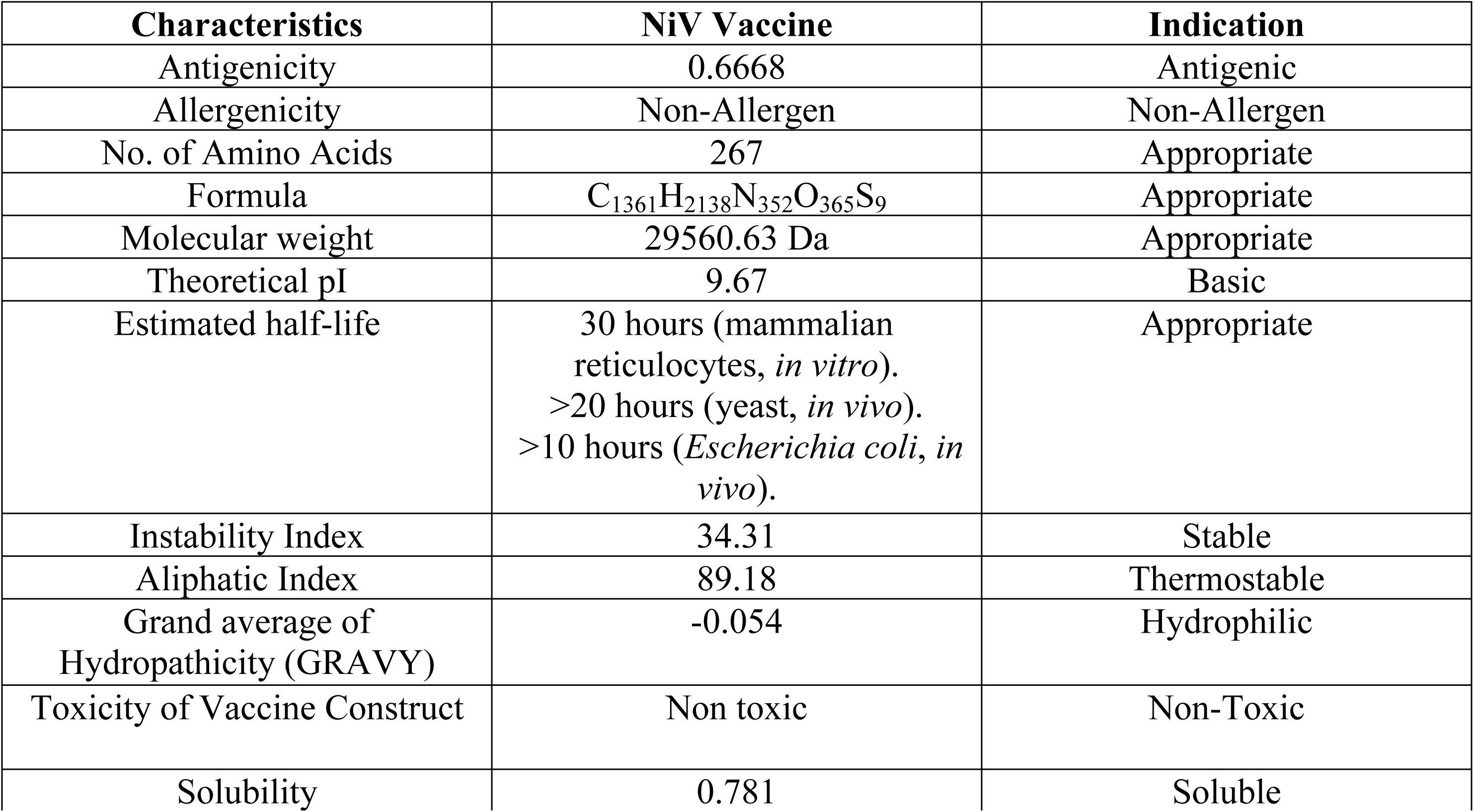
Predicted physicochemical attributes of the proposed multi-epitope vaccine.

Vaccine design contains 267 amino acids and is estimated to have a molecular weight of 29.56 kDa. This construct has a theoretical pI of 9.67 that falls into the basic category. Approximated half-life of the construct in mammalian reticulocytes would be about 30 hours in yeast, more than 20 hours, and in Escherichia coli, more than 10 hours. The modeled instability index points to instability of the proteins. The negative GRAVY of −0.054 indicates that the designs of the vaccines are hydrophilic, i.e., they interact well with surrounding water molecules. To sum up, the additional physicochemical characteristics of the proposed vaccine candidates align with expectations, proving their potential as effective candidates for vaccination.

### Design, refinement, and structural verification of 2-dimensional and 3-dimensional conformations

To determine the structures of the proposed vaccines in a realistic scenario, the structure of each 267-amino acid peptide was predicted in terms of secondary and tertiary structures. The secondary structure prediction performed at the PSIPRED server displays that the vaccine constructs resemble predominantly a variety of helical elements, a small number of β-strands, and an overall trend of coiled structures (see **Fig 4A, B**). In the NiV_1 construct, the structural composition can be estimated at 25.09% alpha-helix, 54.31% random coils, and 20.60% beta-strands.

**Fig 4.**
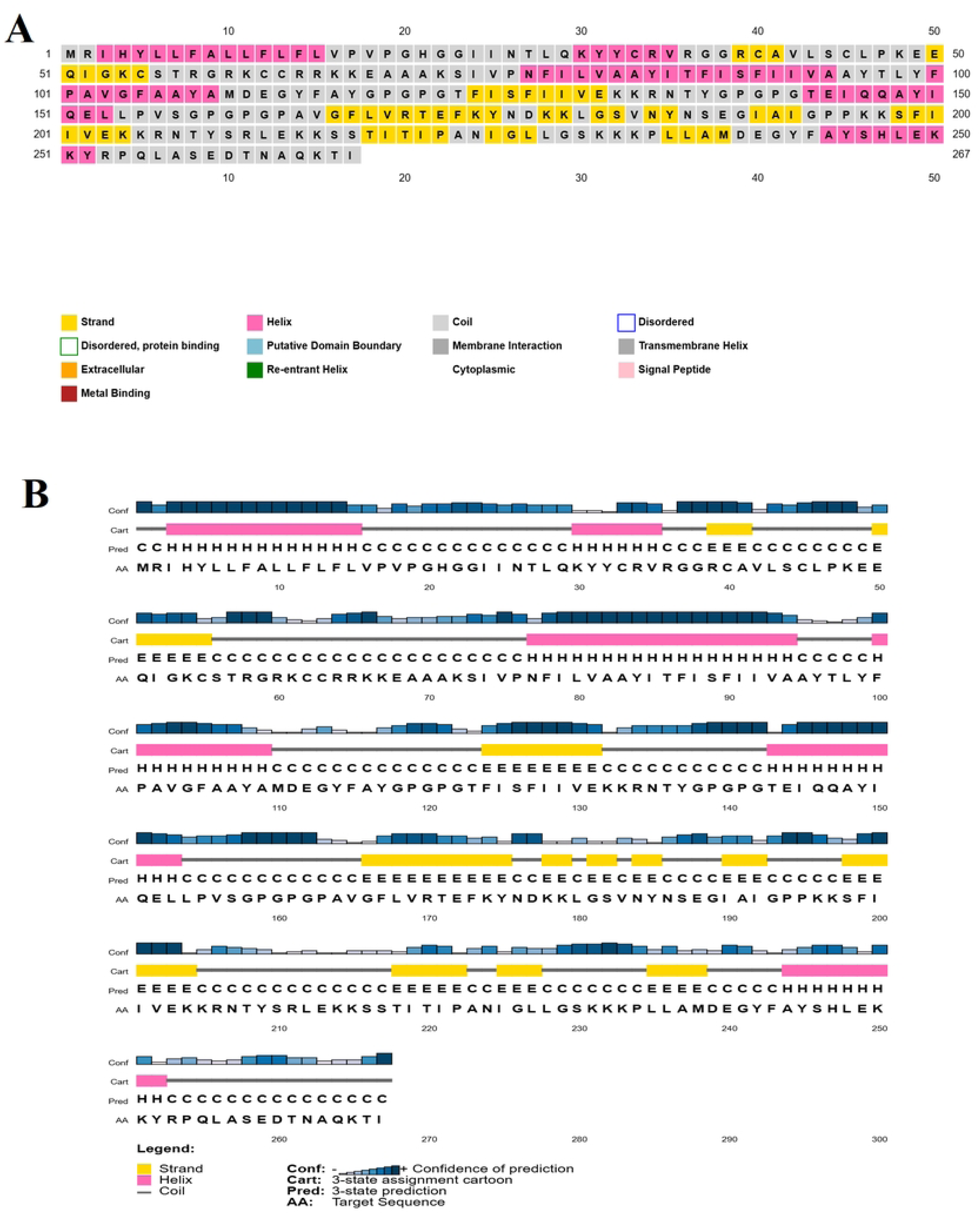
Predicted secondary structure of the designed vaccine construct. **(A)** The figure represents α-helix (pink), β-strand (yellow), and random coils (gray); **(B)** Predicted secondary structure of the vaccine revealed 25.09% α-helix, 20.60% β-strands, and 54.31% random coil regions.

An ideal vaccine candidate needs to have a stable and precise three-dimensional (3D) structure to interact well with the immune receptor proteins of the host organism. The eventual 3D construction of the vaccine went through hard tests of validation and polishing. Through the GalaxyWEB platform, the best structural model was located and later optimized using the Galaxy Refine tool based on the minimum galaxy energy value. Ramachandran plot investigation also added further support to structural validation, which confirmed that in the case of NiV_1 construct, 91.09% of residues were in favored regions, 7.2% in allowed regions, and just 0.9% in disallowed areas in **Fig 5A**. Additional validation of the model reliability was done with the ProSA-web tool to evaluate the structural stability and any possible errors in the model through comparison with known protein structures that have been solved experimentally using NMR and X-ray crystallography in the form of a z-score. The ERRAT server analysis provided the overall quality factor of 87.22, and the ProSA-web tool provided the z-score of - 2.39, which allows concluding that this is a well-validated structure, see **(Fig 5C and 5B**). Together, ERRAT and ProSA-web showed that the protein model of the improved vaccine had a high structural score quality, as shown in **Fig 7A**.

**Fig 5.**
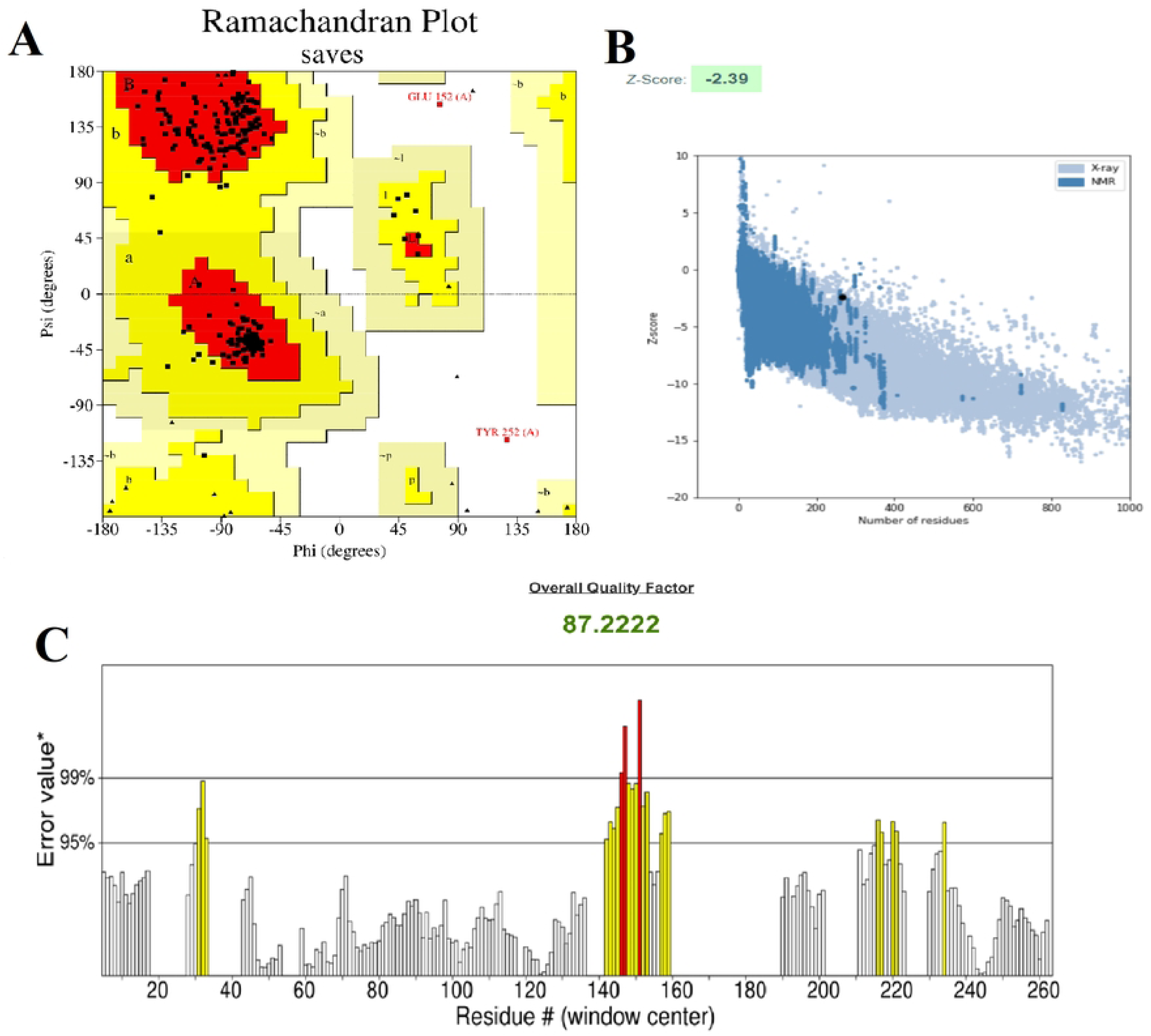
Structure validation and quality assessment for NiV_1 vaccine. (**A**) The Ramachandran plot indicates that for NiV_1, 91.9% of the residues fall within the favored areas, 7.2% in the allowed regions, and 0.9% in the disallowed areas; (**B**) Validation of quality assessment for the refined NiV_1 vaccine by ProSA-web displays a Z-score of −2.39; (**C**) ERRAT analysis validated the refined structure as a high-quality model with a score of 87.22.

### Assessment of conformational B-cell epitopes

The ElliPro database was applied to predict conformational B-cell epitopes on the intended vaccination construct. The analysis revealed the presence of 3 discontinuous epitopes spread in various domains of the protein, and these epitopes of 128 residues were identified with a score between 0.651 and 0.882. Such epitopes were predominantly found in exposed, loop-containing, and flexible regions typically favorable to antibody binding. **Fig 6** illustrates their structural position and the availability of their potential function in the structure’s overall antigenicity.

**Fig 6.**
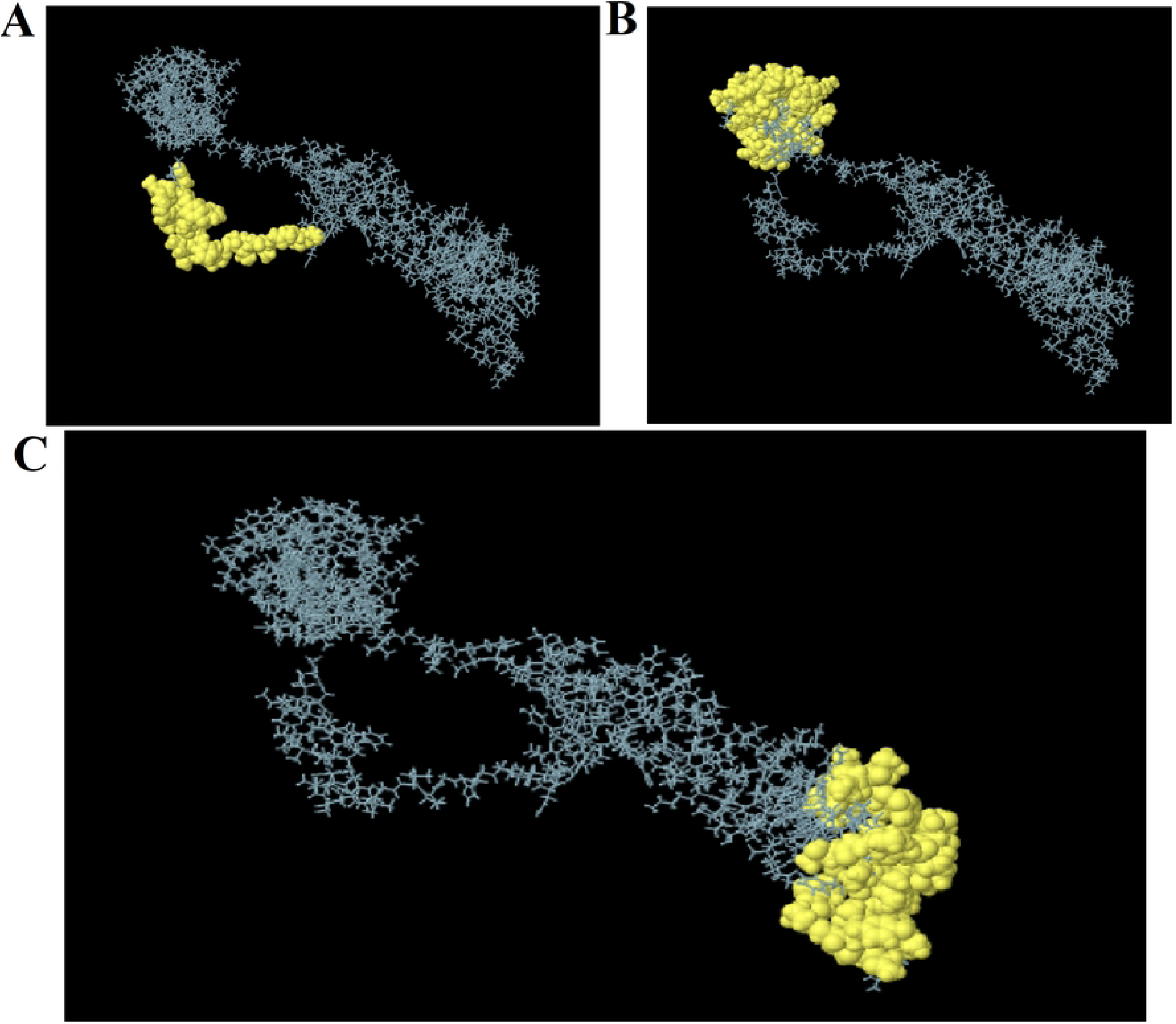
Predicted conformational B-cell epitopes of the vaccine construct. The 3D structure of discontinuous epitopes is marked in yellow, and the non-epitope regions are gray. **(A)** 24 residues with a score of 0.882; **(B)** 65 residues with a score of 0.732; **(C)** 39 residues with a score of 0.651.

### Molecular docking assessment with Toll-like receptor 4 (TLR-4)

The binding affinity between the receptor (TLR-4) and the ligand (refined vaccine) was assessed using the ClusPro 2.0 server. This server produced thirty conformational complexes; each had its binding energy score. **In the S5 Table**, the central and the lowest energy values per cluster are shown. The structure with the lowest binding energy was chosen as the preferred one. Cluster-5, with a binding energy of −1640.9 kcal/mol, was selected for further analysis. The docking of the TLR-4 receptor with the vaccine construct according to the VR docking complex is depicted in **Fig 7B**.

**Fig 7.**
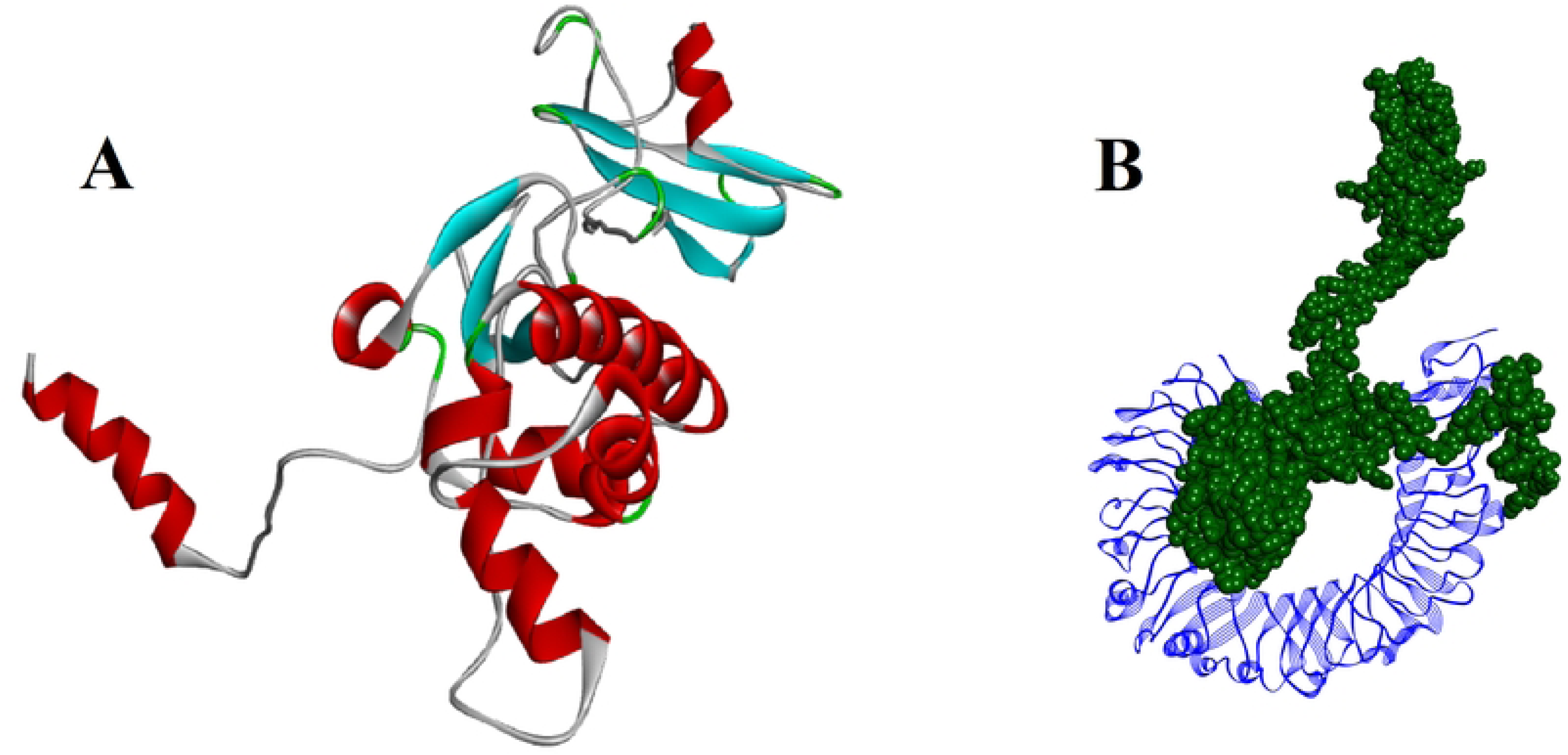
3D structure and molecular docking interactions of NiV_1 vaccine design with human TLR-4. **(A)** Refined 3D structure of the vaccine construct; **(B)** More viable docking interaction, blue indicates the TLR-4 receptor, and green shows the vaccine construct.

### Analysis of the molecular dynamics simulation of the vaccine complex

MDS is an effective method of analyzing the conformational stability of molecules and atoms through the simulation of cell interactions at the atomic scale. Its outstanding strength is the capacity to bring out the integrity of the ligand in a particular protein macromolecule. In the current research, a 100-nanosecond MD simulation was conducted to examine the structural orientation of the chosen vaccine complex. The objective was to establish the ligand potential to bind with the protein, especially in the active site cavity. The results generated during the simulation were evaluated based on parameters like RMSD, RMSF, radius of gyration (Rg), and intramolecular hydrogen bonds (Intra HB).

### Root mean square deviation of vaccine construct

The root mean square deviation (RMSD) in the vaccine construct was calculated to elicit the average movement of chosen atoms compared to a frame of reference. During the run, the greatest fluctuation was 21.113 Å, the least movement was 2.103 Å, and the average movement was 14.124 Å (**Fig 8A**). At 70 ns, a minor fluctuation was observed, which shows that the constructs have a stable structural dynamic.

**Fig 8.**
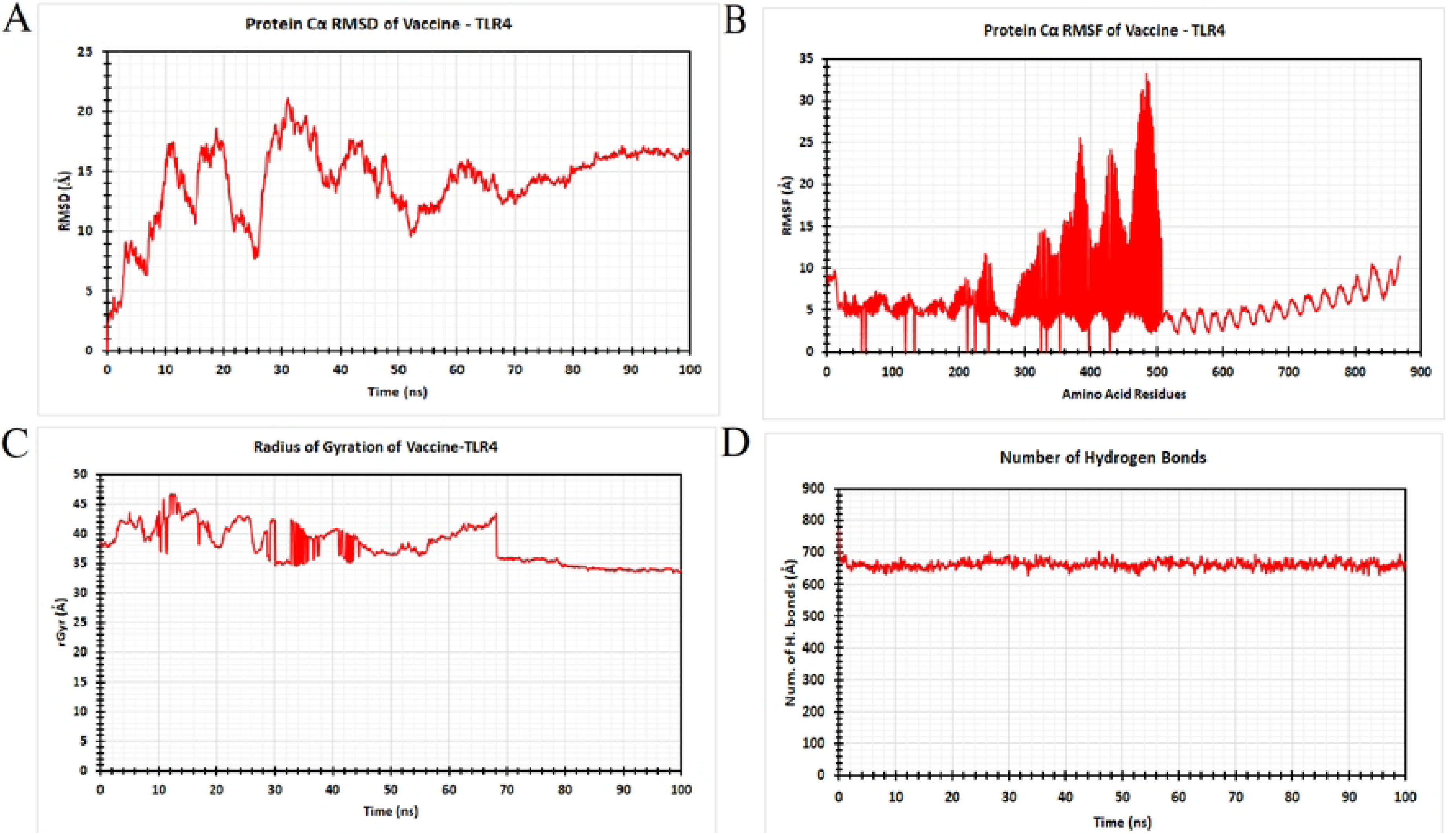
Molecular dynamics simulations of TLR-4–vaccine complexes were performed for 100 ns using the Schrödinger software. **(A)** RMSD; **(B)** RMSF; **(C)** Rg; **(D)** HB interaction analysis with the TLR-4 receptor and vaccine complex.

### Root mean square fluctuation of vaccine construct

The Root mean square fluctuation (RMSF) analysis was used to gain more information about structural stability. The RMSF plot of the vaccine receptor (VR) complex showed that the regions with the most variation were residues 460-500 and 360-400, with a value of 33.332 and 25.392 Angstroms, respectively. Outside these areas, the VR complex had relatively good regularity with slight fluctuations (**Fig 8B**). The findings of these observations reflect the significance of RMSF analysis as a tool to better comprehend structural dynamics and design optimization of the NiV vaccine candidate, which will eventually boost its immunogenicity and protective capability.

### Radius of gyration of vaccine construct

In **Fig 8C**, the construct’s structural compactness was assessed using the radius of gyration (Rg). The Rg profile showed higher variances between 5–68 ns, with an average value of 38.03 Å. The construct’s overall low Rg value suggested a tightly packed structure, which is a characteristic of α/β-proteins.

### Number of hydrogen bonds of vaccine construct

A hydrogen bond (HB) analysis further evaluated the structural stability. Their fluctuation distribution between the donor and acceptor atoms was assessed to ascertain the dynamics of HBs during the 100 ns simulation (**Fig 8D**). The construct demonstrated the maximum amount of HBs within 5-100 ns, with an average of 662.93 bonds. This implies that the vaccine construct can achieve dynamic equilibrium to retain its active confirmation via extensive hydrogen bonding.

### Host-specific codon adaptation and *in-silico* cloning

The NiV_1 vaccine construct was selected due to positive results from molecular docking and molecular dynamics simulations. The Vector Builder server optimized codons to determine their potency to be expressed in the *E. coli* system. The optimized codon sequence of the vaccine was made of 804 nucleotides. An ideal expression within the host was evidenced with a codon adaptation index (CAI) of 0.94 above the optimal range of 0.8 and the GC content of more than 51.24% which falls within the 30% to 70% optimal range. To support cloning, XhoI and BamHI restriction sites were introduced at the N- and C-terminal positions of the optimized sequence. Then, this construct was introduced into the pET-28a (+) plasmid using Snap Gene software. **Fig 9** shows a graphical representation of the cloning procedure and the vaccine construct NiV_1 insertion into a vector.

**Fig 9.**
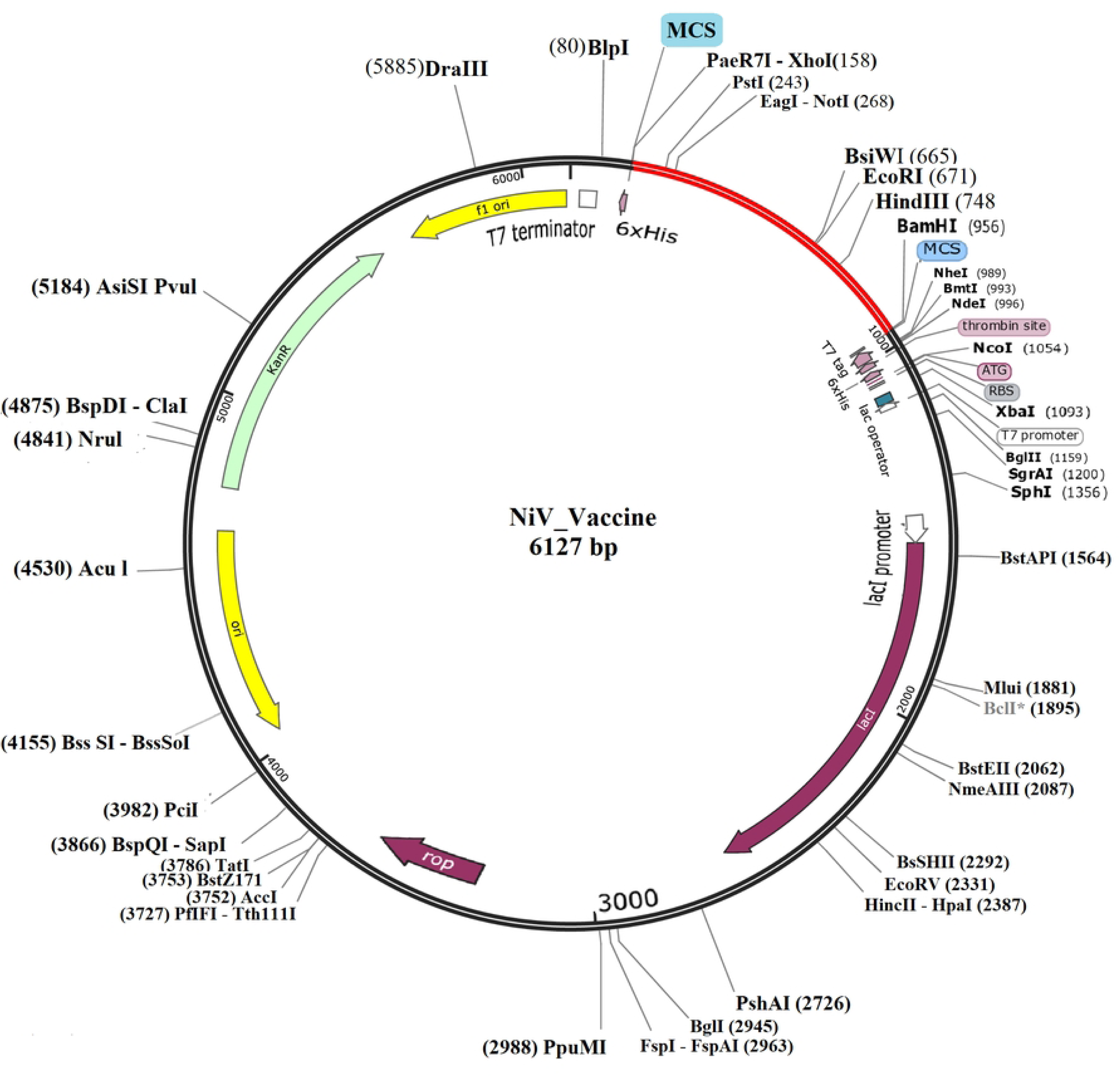
*In silico* vaccine construction of the NiV_1 in the pET28a+ expression system utilizing Snap Gene. The red arrow indicates the inserted piece of DNA being cloned, and the rest of the fragment represents the backbone of the vectors.

### *In silico* simulation of immunological responses

The C-ImmSim web platform was used to simulate the effect that the proposed vaccine candidate should have on the immune response. The findings indicate that the construct could induce a high immune response. Within the first five days of administration, the antigens were found at about 700000 antigens/mL and were quickly eliminated from the host system. After 28 days of vaccination, when the antigen levels reduced to almost 500,000 antigens/mL, there was a significant increase in the production of primary antibodies; the levels of IgM and IgG increased to above 9000 units, and after 56 days, they reduced to 120000 units. Subsequently, the antigen-antibody complex levels declined, but they stagnated at around 40000 on day 100, which implies a long-lasting, stable immune memory (**Fig 10A**). After vaccination, the overall B cell levels exceeded 450 cells/mm³, and the memory B cell and IgM numbers were also maintained during the simulation (**Fig 10B**). The B cells in the plasma reached peak on the 30-60 days and exceeded a value of 60 cells/mm³, along with more IgM and IgG1. Notably, the results shown in **Fig 10C** support an exaggerated increase in the number of plasma-producing B cells, which reveals their importance in pathogen clearance.

**Fig 10.**
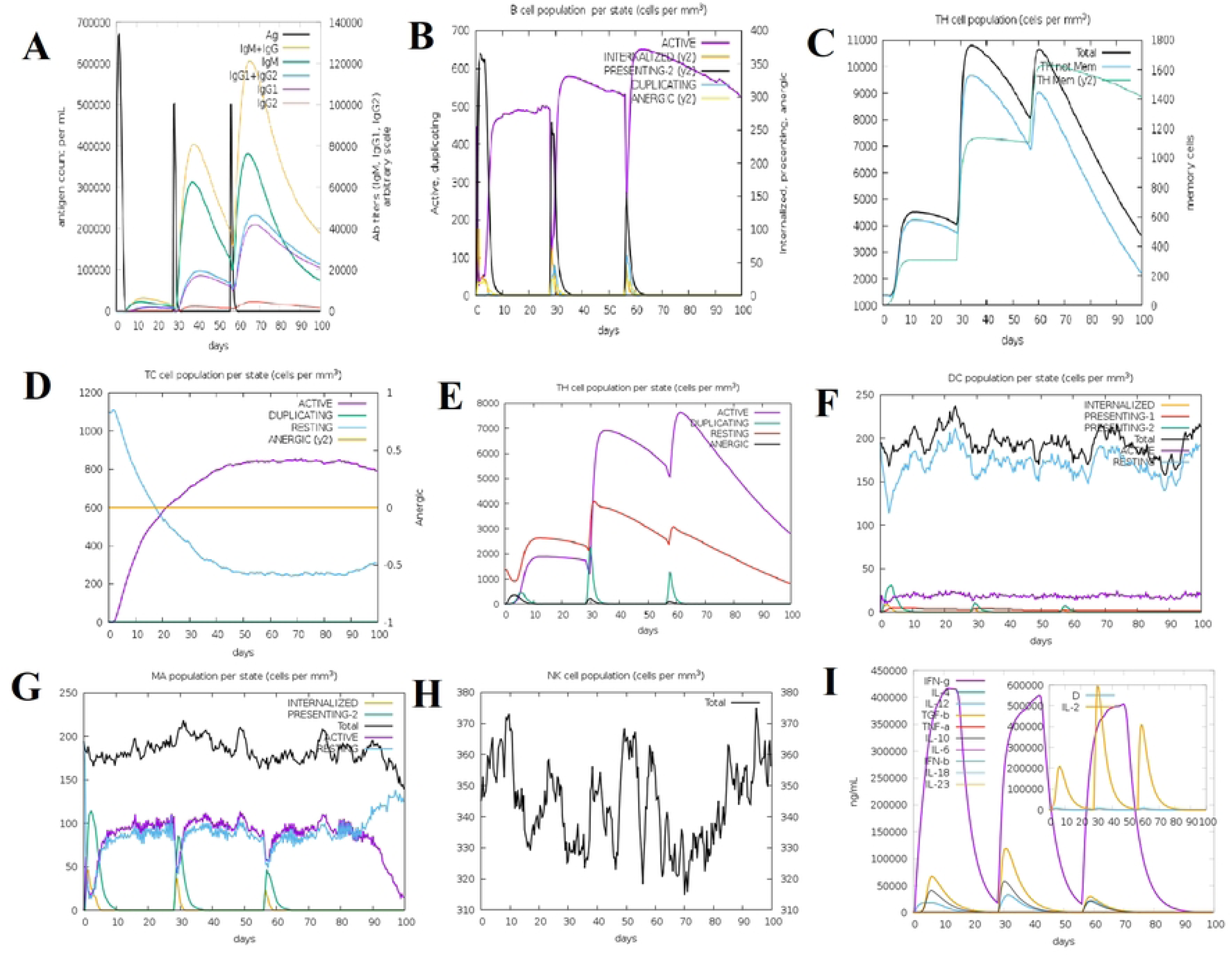
Immune response simulation by the c-ImmSim server following a single administration of the NiV_1 vaccine. **(A)** Generation of immunoglobulins and B-cell isotypes following exposure to the antigen; **(B)** count of active B-cells in each state; **(C)** quantity of plasma B-lymphocytes and their isotypes in each state; **(D)** population of helper T-cells; **(E)** condition of the helper T-cell population during subsequent immune responses; **(F)** quantity of cytotoxic T-cells in antigen-exposed states; **(G)** activity of macrophage populations during three sequential immune responses; **(H)** dendritic cell population in each state; and **(I)** production of cytokines and interleukins along with the Simpson index of the immune response.

The ability of the vaccine to trigger the release of cytokines was also tested, and it showed a large induction of interferon-gamma and interleukin-6 with the highest concentration of 400,000 ng/mL and 600,000 ng/mL, respectively (**Fig 10D**). Further, the number of helper T cells (Th) and cytotoxic T cells (Tc) peaked greatly following vaccination, as indicated in **Fig 10E and Fig 10F**. In its entirety, C-ImmSim analysis suggests that the NiV_1 vaccine candidate can potentially induce both an effective cellular and humoral immune response. Nevertheless, additional experimental confirmation in the laboratory is required to confirm its efficacy in eliciting adaptive immunity against NiV_1.

## Discussion

Vaccination is an important and remarkable achievement in the pharmacology history. A great number of lives have been saved by vaccination procedures (56). According to the World Health Organization, around 2-3 million deaths of people are prevented each year through vaccination. Additionally, vaccination offers economic benefits by reducing medical spending for disease and hospitalizations by over $500 billion (57). The fatal diseases, including Ebola, SARS-CoV-2, Zika, and Nipah virus, have underscored the major risks to public health caused by newly discovered viral infections (58). Vaccination boosts the host defense system by suppressing various fatal diseases. For this reason, immunological response protects both individuals and public health. Recent developments in immuno-informatics mixed with bioinformatics and other computational tools have greatly optimized the vaccine design process, reduced expenses, and lowered time while increasing the reliability of peptide-based vaccine strategies (59). Several research studies have utilized immuno-informatics approaches to design vaccines against numerous viral pathogens, including the SARS-CoV-2, Lassa virus, herpes simplex virus, chikungunya virus, influenza virus, yellow fever virus, and dengue virus (60,61). Nipah virus (NiV) reflects an important emerging threat with the potential to cause pandemics. Currently, no therapeutics or vaccines are approved against this virus for humans or livestock use (62). Nipah, an RNA virus (NiV), reveals a strong inclination for spontaneous mutations. These alterations can accelerate viral escape from immune selection if vaccines are directed against a small number of antigens. Thus, developing a poly-epitope vaccine targeting different types of antigens is essential for designing vaccines for extensive protection against the Nipah virus and reducing the risk of immune evasion due to genetic variation. Hendra virus (HeV) soluble G glycoprotein (sG) is the most thoroughly investigated among NiV vaccine candidates. Targeting the F and G envelope glycoproteins is a key strategy in NiV vaccine research, as these proteins represent the primary target sites engaged by neutralizing antibodies (63)(64).

This study aims to develop an epitope-based vaccine targeting different antigenic structural peptides of the Nipah virus (NiV). Our investigation represents the first design of a distinct vaccine construct with adjuvants and performs its comparative evaluation through bioinformatics analyses. This approach employs different types of in-silico and immuno-informatics techniques. It improves the ability to stimulate a strong and long-lasting immune response. In the present study, antigenicity-specific analysis was performed with protein sequences obtained from the Uniprot database. After this investigation, these sequences underwent epitope prediction. This aimed to induce B-cell (humoral) and T-cell (cellular) mediated immunological effect using different immuno-informatics methods.

Cytotoxic T cells identify antigens presented on body cells. Cytotoxic T cells (CD8+) identify foreign antigens like endogenous peptide fragments derived from pathogens’ proteins are displayed on infected body cells through MHC-I molecules. On the other hand, antigen-presenting cells (APCs) like macrophages, B cells or dendritic cells are recognized by CD4+ Helper T cells via class-II MHC molecules. Helper T cell is activated by the interaction between antigen and APCs provides signals to enhance B cell proliferation and antibody production (65). The identification of potential CTL vaccine candidate epitopes involved further evaluation of their conservancy, allergenicity, antigenicity, toxicity, and immunogenicity (for CTL epitopes), while IFN-gamma induction analysis was performed only for HTL epitopes. T-cell epitopes can bind with multiple HLA-I/II alleles. Choosing these epitopes helps to cover a broader population and improve the potential for an effective immunological reaction. The vaccine was designed with 5 HTL, 13 CTL, and 16 B-cell epitopes for the generation of initial and memory immune responses. The selected T-lymphocyte epitopes analysis covers around 97.94% of MHC alleles worldwide. The analyzed epitopes were connected through different linkers, namely GPGPG, AAY, KK, and EAAAK. Linkers play a vital role in ensuring vaccine structural stability, interdomain interactions, and maintaining the overall functionality (66). Selection of these linkers was mainly based on their potentiality. The EAAAK linker was employed to minimize interference between the adjuvant and its receptors while ensuring structural rigidity. The EAAAK linker is used to attach MHC-II epitopes. B-cell immunogenicity and pH stability are maintained by the KK linkers while the AAY linkers join CTL epitopes through a proteasomal cleavage site. Moreover, KK linkers help to maintain spacing between epitopes. The GPGPG linker is used to effectively stimulate a helper T-cell response, enhancing immunogenicity for the vaccine, and connecting the MHC-I or HTL epitopes (67,68).

A peptide vaccine’s potential is mainly determined by the selected adjuvant. Adjuvants are one kind of substance that is incorporated into vaccines for enhancing immune stimulation, ensuring stronger and longer-lasting protection (69). The innate immune defense system is activated by an antimicrobial peptide known as Beta-defensin. Evaluation of the peptide was conducted to determine a suitable adjuvant. Research evidence has revealed the function of Beta-defensin as an immune regulator against diverse pathogens (70). In addition, β-defensin enhances adaptive immunity through the recruitment of naive T cells and immature dendritic cells (DCs) to infection sites (71). Compared to traditional adjuvants, Beta-defensin 3 provides unique immune-enhancing benefits. It improves the vaccine’s potential for mucosal-targeted vaccine development. Analysis confirmed that the developed vaccine constructs validated as immunogenic and allergenicity and toxicity free. It is essential to evaluate the physicochemical traits of the engineered vaccine construct to ensure its safety and potency as a multi-epitope vaccine (72). The ProtParam server was utilized to estimate these properties. The analysis demonstrated that the vaccine design is composed of approximately 267 amino acids and exhibits a molecular weight of 29.56 kDa. Additionally, these findings showed that the vaccine construct is stable, soluble, hydrophilic, and thermostable, leading to prolonged stability half-life. PSIPRED-based secondary structure prediction revealed that the construct is mainly helical with a few β-strands and coiled regions. Tertiary structure of the vaccine construct was predicted with the GalaxyWEB platform and further refined with the Galaxy Refine tool. Further, Ramachandran plot analysis, ERRAT, and ProSA-web server were used for structural validation of the 3D construct with a confirmed Z score.

The vaccine activates the immune system via binding to immune receptors, allowing it to recognize and respond to foreign particles (73). TLR-4, a transmembrane receptor known as toll-like receptor 4, regulates natural and adaptive immune responses towards microbial infections and cellular damage through the production of IFN (74). In this study, molecular docking analysis and molecular dynamics (MD) simulations were performed with TLR-4 to investigate the stability and binding affinity of the vaccine construct. This study identified the method by which vaccines interact with immune cells to elicit a strong immunological response in the host. The most advantageous efficacy was demonstrated by the TLR4-NiV_1 combination with the lowest binding energy. The Vector Builder tool was utilized in this study to optimize codons and improve expression efficiency in the E. coli system. The vaccine design had a codon adaptation index (CAI) of 0.94 and a GC content of 51.24%, indicating a greater potential for gene expression in E. coli. For in silico cloning, restriction sites were added, and the construct was subsequently added to the pET-28a (+) plasmid. The multi-epitope vaccination demonstrated strong immunological stimulation in the human body, which was subsequently investigated using the C-ImmSim server. Following the first administration of a single vaccine, a significant increase in the secretion of IgM and IgG antibodies, IFN-gamma, IL-6, B and T-cells was observed. These results revealed the ability of the design vaccine construct to induce a protective immune response against NiV (75). Even though the computational method employed to develop a vaccine against NiV follows high standards, limitations still require further research. Although the *in-silico* pipeline provided a logical rationale in the design of vaccines and showed promising immunogenic potential, these outcomes cannot be considered final. Strict post-analysis and epitope filtering criteria made the vaccine a more promising candidate against the Nipah virus. Still, the results remain tentative and must be confirmed through test tube (*in vitro*) and animal (*in vivo*) models. This experimental testing will be necessary to validate its immunogenicity, safety, and translational feasibility before clinical use.

## Conclusion

The continual outbreak of Nipah virus (NiV) infections highlights the pressing need for an efficacious and broad-spectrum multi-epitope vaccine without commercial antiviral drugs. Immunoinformatics and bioinformatics methodologies were also used in this investigation to generate vaccine designs that target NiV fusion and glycoproteins. These constructions were discovered to possess desired properties such as antigenicity, immunogenic potential, non-allergenic behavior, and safety, in addition to meeting severe physicochemical parameters. Structural validation employing molecular docking and dynamics simulation indicated that the vaccination was stable and had a high binding affinity with a toll-like receptor 4 (TLR-4) that strengthened the vaccine. Additionally, the wide range of protection is made possible by the fact that the chosen epitopes are conserved across different NiV strains. The combined results support the vaccine’s function in promoting the body’s innate and adaptive defenses against NiV. Since these results are merely predictive, experimental validation is required to ensure immunological efficacy, accuracy, and safety.

